# OT-Mation: an open-source code for parsing CSV files into python scripts for control of OT-2 liquid handling robotics

**DOI:** 10.1101/2025.01.20.633898

**Authors:** Alex Laverick, Katherine Convey, Catherine Harrison, Jenny Tomlinson, Jem Stach, Thomas P. Howard

## Abstract

OT-Mation is an open-source Python script designed to automate the programming of OT-2 liquid handling robots, making combinatorial experiments more accessible to researchers. By parsing user-defined CSV files containing information on labware, reagents, pipettes, and experimental design, OT-Mation generates a bespoke Python script compatible with the OT-2 system. OT-Mation enhances reproducibility, reduces human error, and streamlines workflows, making it a valuable addition to any laboratory utilising OT-2 robotics for liquid handling. While OT-Mation can be used for setting up any type of experiment on the OT-2, its real utility lies in making the connection between multifactorial experimental design software outputs (i.e. Design of Experiments arrays) and liquid-handling robot executable code. As such, OT-Mation helps bridge the gap between code-based flexibility and user-friendly operation, allowing researchers with limited programming skills to design and execute complex experiments efficiently.

## INTRODUCTION

Automation of small-scale liquid handling is gaining traction in life sciences, allowing researchers to conduct high throughput or combinatorial experiments with reduced human error, greater accuracy and with traceable workflows^1^. A combination of low-cost and flexibility makes their incorporation into existing workflows an attractive option for many researchers^2–4^. There is also recognition that a large proportion of scientific research cannot easily be replicated, and that part of the problem may be the quality and consistency of protocol reporting^5–7^. The explicit and shareable protocols used by liquid-handling robotics may therefore be a useful tool to improve data quality and reproducibility^1^. Another contributing factor may be the use of the one-factor-at-a-time (OFAT) experimental approach. This persists despite well-documented drawbacks including (i) intuition, narrative and confirmation biases and (ii) sensitivity to unknown confounding factors^8, 9^. Automated liquid handling can increase the combinations of experiments that can be handled by an individual researcher, facilitating better designed experiments with more robust outcomes.

Combining automated liquid handling robotics with statistical Design of Experiments (DoE) offers a powerful, iterative approach to understanding and optimising complex biological systems. DoE provides a robust framework for experimental design, minimising experimental effort whilst maximising knowledge gained. Compared to the traditional OFAT approach, DoE separates real effects from experimental noise with far greater efficiency and uncovers multifactorial interactions^8, 9^. When coupled with modern approaches to data analysis, it provides predictive models for experimental optimisation. Synthetic biology already applies DoE in a range of settings^10^, including understanding and manipulating metabolic pathways^11–13^, engineering biosensors^14^, and optimising cell-free systems^15–17^.

Despite the apparent synergy between DoE and automated liquid handling, the ability to couple these two approaches is not straightforward. Protocol development for liquid handling robotics is typically via either a Graphical User Interface (GUI) or a computer programming language. Creating a simple protocol through a GUI is quick and easy and offers an excellent entry into the use of robotics for new users. Implementing and editing more complex procedures is difficult and may take more time than running the protocol^15, 18^. By contrast, some liquid handling platforms permit direct programming of the robotics, allowing users to develop more bespoke protocols. The OT-2 system (opentrons.com) operates through Python scripts providing flexibility and control. Programming in Python, however, requires skills that many laboratory-focused researchers are not confident in^19^. Furthermore, while multifactorial experiments can be set up using code, critically DoE is a sequential optimisation process that frequently requires fold-over or augmentation of experimental designs with new combinations of factor settings^9^, requiring a new script for each sequential experiment. Finally, both methods require human intervention to transfer experimental arrays into the liquid handling protocols, which may also introduce errors into the process. Neither GUI nor direct programming is therefore easily adapted for the multifactorial experiments required by DoE.

Here we describe OT-Mation, a Python script that automates and simplifies the programming steps of the OT-2 without sacrificing the utility of code-based development. OT-Mation generates a bespoke Python script for control of the OT-2 from four user defined comma-separated value (CSV) format files. These files provide information on the labware on the deck of the robot, the reagents required for the experiment, pipetting hardware, and the experimental design parameters. In practise, only the experimental parameters file needs modification between experiments, and its format aligns with experimental arrays from statistical design software, eliminating the need for manual data entry into a GUI or Python code.

## RESULTS AND DISCUSSION

### Design considerations

OT-Mation was designed with flexibility and iteration in mind. The script is capable of supporting complex, iterative workflows that utilises a broad variety of commands and enables users to specify settings for each reagent to give the user control over their experiments. Our primary motivation was to support sequential multifactorial experimentation as required when using DoE, however, the versatility of OT-Mation makes it suitable for a variety of experimental procedures (e.g. serial dilutions or setting up PCR). In a statistically designed experimental array there are multiple experimental runs in which the concentrations (or sometimes a categorical selection) of multiple factors are adjusted simultaneously and systematically. When run, OT-Mation generates an output script that once written, remains the same. This allows scripts to be archived alongside experimental data, acting as a record of experiments carried out.

### File structure

OT-Mation parses the data contained within four CSV files to generate an output Python script that is uploaded onto the OT-2 through the Opentrons companion application (Figure 1). The four CSV files are *Experimental Parameters, Labware Inventory, Stock Inventory* and *Pipette Settings* (Figure 2). OT-Mation calculates the pipetting actions for the robot to complete the experiment without requiring the user to write any code.

**Figure 1.**
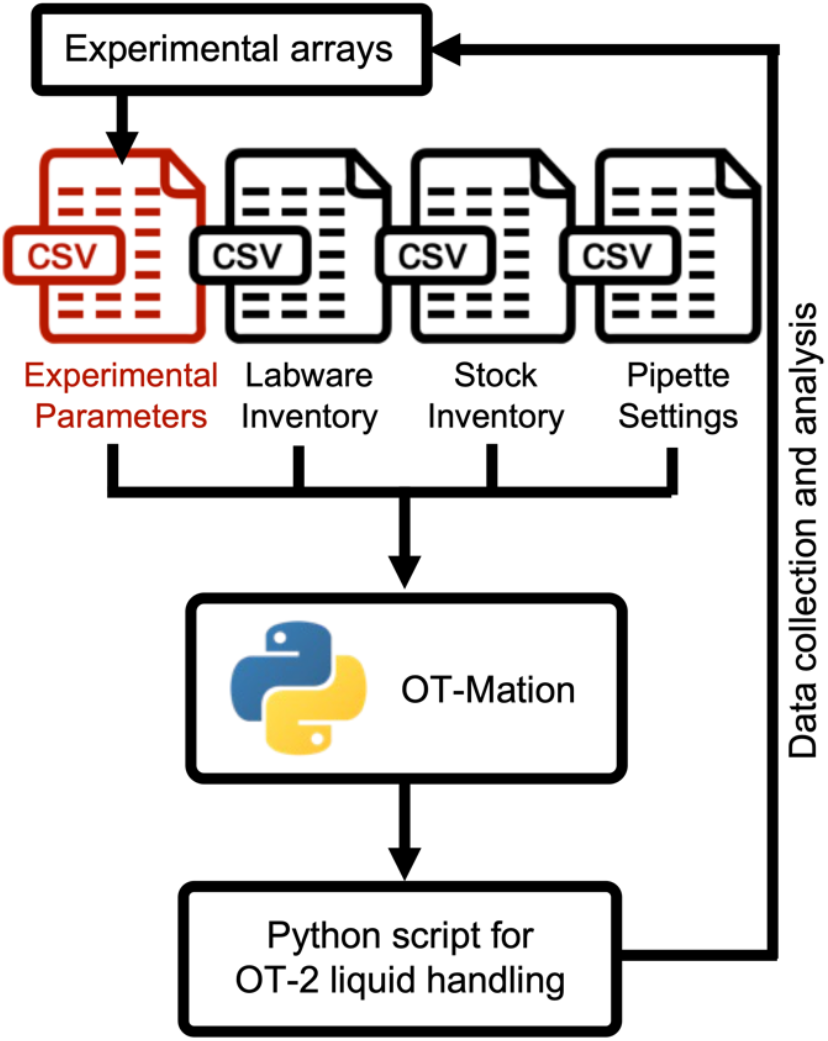
OT-Mation workflow. OT-Mation was designed to support the iterative, sequential workflow required in statistical Design of Experiments but can be applied in any iterative Design-Build-Test-Learn cycle. In a cyclic experimental process, after results from an experiment have been analysed, new factor combinations or parameters are chosen. Using OT-Mation, only the *Experimental Parameters* CSV file (red) needs to be changed to generate a new protocol - and this is compatible with the output format of many statistical packages removing the need for human intervention in transferring data.

**Figure 2.**
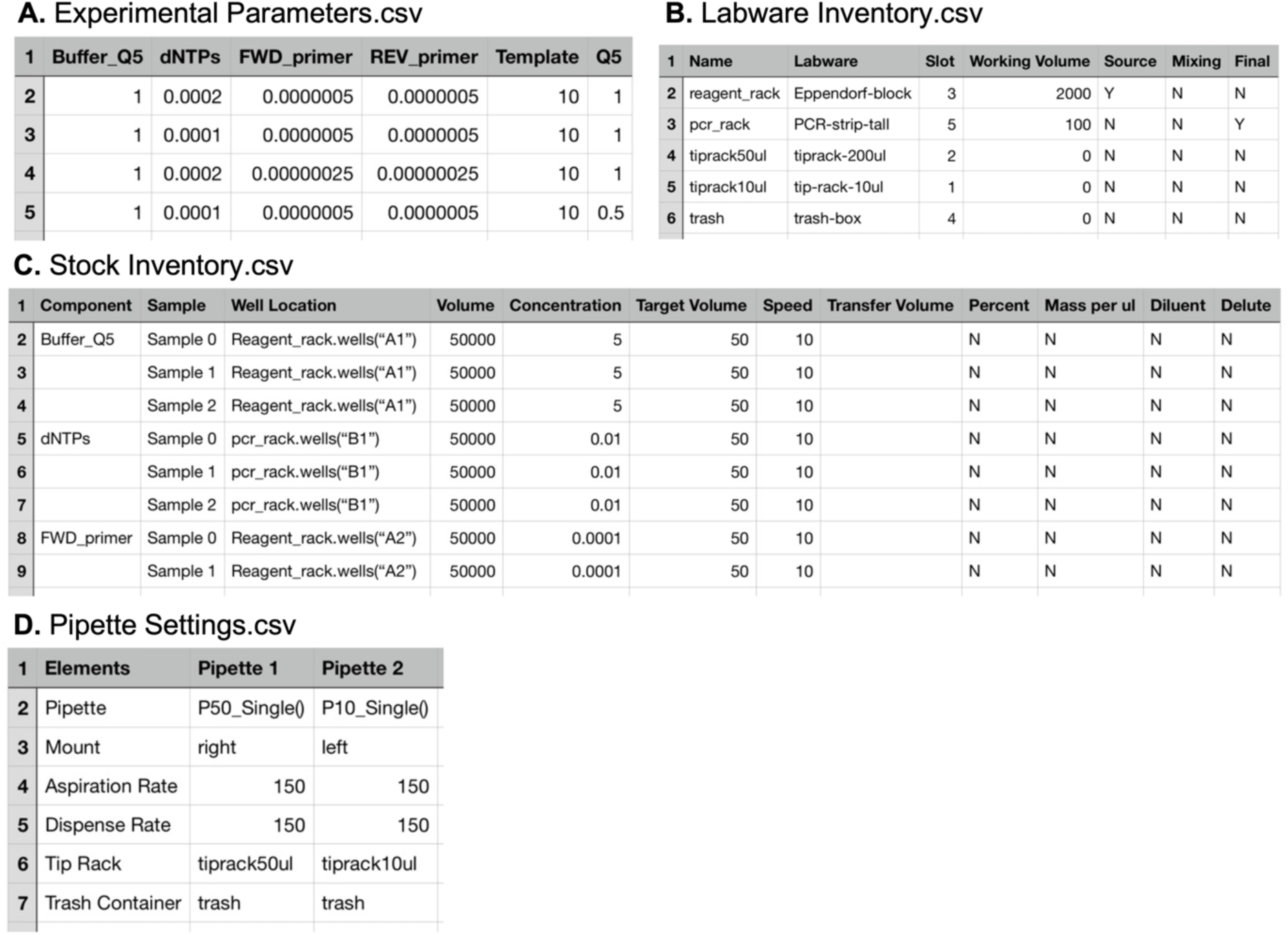
Layout of the four user defined CSV Files central to OT-Mation. **A**. Experimental Parameters. **B**. Labware inventory. **C**. Stock Inventory. **D**. Pipette Settings.

#### Experimental Parameters File

The Experimental Parameters file is the primary route by which experimental designs are parsed into OT-Mation. Between experimental iterations, this is the only file that needs to be altered. Each row represents a sample being assembled and each column a reagent (Figure 2A). Each row represents an experimental run as identified in the experimental array, the first row is for the headers, the second row is Sample 0, the third row is Sample 1, and so forth. In each cell, the desired final concentration of the reagent in the sample is entered. If 0 is entered, the reagent is skipped. OT-Mation calculates the volume of Diluent required to reach the target volume of the sample. If the number of rows within Stock Inventory exceeds the rows of Experimental parameters, OT-Mation will produce an error.

#### Labware Inventory File

All labware to be used in the OT-2 is specified in the Labware Inventory CSV file (Figure 2B). Each row specifies one piece of labware, and all information about that labware is found within seven columns: *Name, Labware, Slot, Working Volume, Source, Mixing* and *Final*. The *Name* column identifies the name used within the Python script, while the *Labware* column specifies the labware according to the Opentrons labware library. *Slot* refers to which working position the labware will be in, as the OT-2 has 11 fixed labware slots. *Working Volume* specifies the maximum volume each piece of labware can hold, given in microlitres. Labware that is not used for mixing and storing samples, such as tip racks and trash, can be assigned values of 0. A piece of labware is designated as *Source, Mixing* or *Final* by marking the cells in the last three columns with a Y (yes) or N (no). Labware designated as a *Source* will only be used as a location to aspirate reagents from, *Mixing* identifies if the labware is acting as an intermediary mixing platform and *Final* identifies labware as the final destination for samples. The information in *Source, Mixing* and *Final* are combined with data from the Experimental Parameters CSV file to write the protocol and determine how reagents are handled.

#### Stock Inventory File

The stock inventory is recorded across columns with each reagent represented across multiple rows according to the concentration of the reagent in each sample (Figure 2C). Entries in the *Component* column are parsed by OT-Mation and designated as keys for the dictionary used to store experimental sample information. If a cell is blank, the components designation is considered the same as the previous named entry. *Sample* refers to the assembly of each sample by the OT-2. This starts at 0 as this is the initial value for lists and counts in Python. *Well Location* refers to where the reagent stock is stored on the OT-2 work surface and what piece of labware it is stored in. *Volume* indicates the volume of the stock available, in microlitres. *Concentration* indicates the concentration of the stock solution. The concentration can be recorded as percentage, mass per microlitre or mols. *Target Volume* refers to the final volume of the sample in microlitres. Information recorded in the *Speed* column sets a default value for both aspirating and dispensing rate in microlitres per second. Values entered here overwrite the default speed in Pipette Settings. *Transfer Volume* is the specific amount of a reagent to be dispensed into the samples. If cells in this column are left blank, OT-Mation will calculate which transfers need to occur to create a sample containing the correct concentration of reagents. Users can specify if a reagent is for diluting a sample or another reagent, or if the reagent needs to be diluted by a diluent before it is used in sample assembly by entering Y or N in *Dilutant* and *Diluent* columns. Reagents are handled by the OT-2 in the order they are listed in the Stock Inventory file, distributing reagent A into all samples before reagent B is used.

#### Pipette Settings File

The Pipette Settings CSV contains the settings for the pipettes used throughout the assembly of the samples (Figure 2D). Two pipettes can be mounted simultaneously in the OT-2. The columns of the pipette setting CSV file represent the two pipettes, and each row specifies their settings, including *Pipette, Mount, Aspirate Rate, Dispense Rate, Tip Rack* and *Trash Container. Pipette* specifies which model of pipette (P10, P50, etc) is being used. This needs to be entered as it appears in the Opentrons API. *Mount* indicates which mount the pipette is affixed to, either left or right. *Aspirate Rate* and *Dispense Rate* sets the aspiration rate and dispense rate of the pipettes respectively, in microlitres per second. If a speed is not specified in the Stock Inventory CSV for a specific reagent, the aspirate and dispense rate in Pipette Settings is used. *Tip Rack* allocates a tip rack for the corresponding pipette. The names entered in these cells must match names given to tip racks in the Labware Inventory. *Trash Container* assigns a location for each pipette to eject tips. OT-Mation automatically calculates which pipette to use based on the volume that needs to be transferred at each step.

### Implementation

OT-Mation runs in the same way as a standard Python script, through the user’s preferred integrated development environment (IDE) or text editor. The four CSV files must be within the same working directory as the OT-Mation script to be read. Alternatively, OT-Mation can be edited to contain file roots to the CSV files of interest. Within the OT-Mation script, there are spaces designated for the file names of the four CSV files, and for the file name of the output script (Figure 3). This is the only time users need to edit the Python script. As the CSV files resides in the same folder as OT-Mation, OT-Mation can be copied into multiple folders with the files of interest to keep work separate^20^.

**Figure 3.**
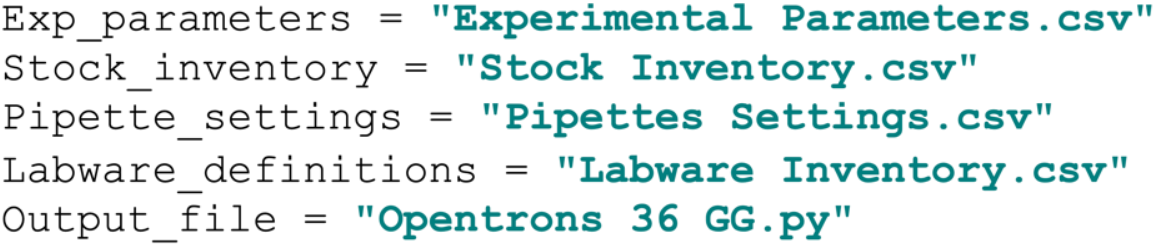
File declarations. OT-Mation File declarations occur within the first five lines of the OT-Mation script meaning that users need only change the coloured text within the quotations marks of the declarations to match the CSV files they wish OT-Mation to read. Users can also specify the name of the output file which will be generated in the same working directory.

OT-Mation reads the file contents in a specific order: Stock Inventory, Experimental Parameters, Pipette Settings and finally Labware Inventory, with Stock Inventory serving as a reference for dictionary assembly. Dictionaries ascribe ‘Keys’ to ‘Values’, for rapid access to CSV data, which is then stored in nested dictionaries. (Figure 4). Nested dictionaries efficiently catalogue data for automated scripts, enabling fast calculations for dilutions, well transfers, reagent stocks, and target concentrations without affecting unrelated data.

**Figure 4.**
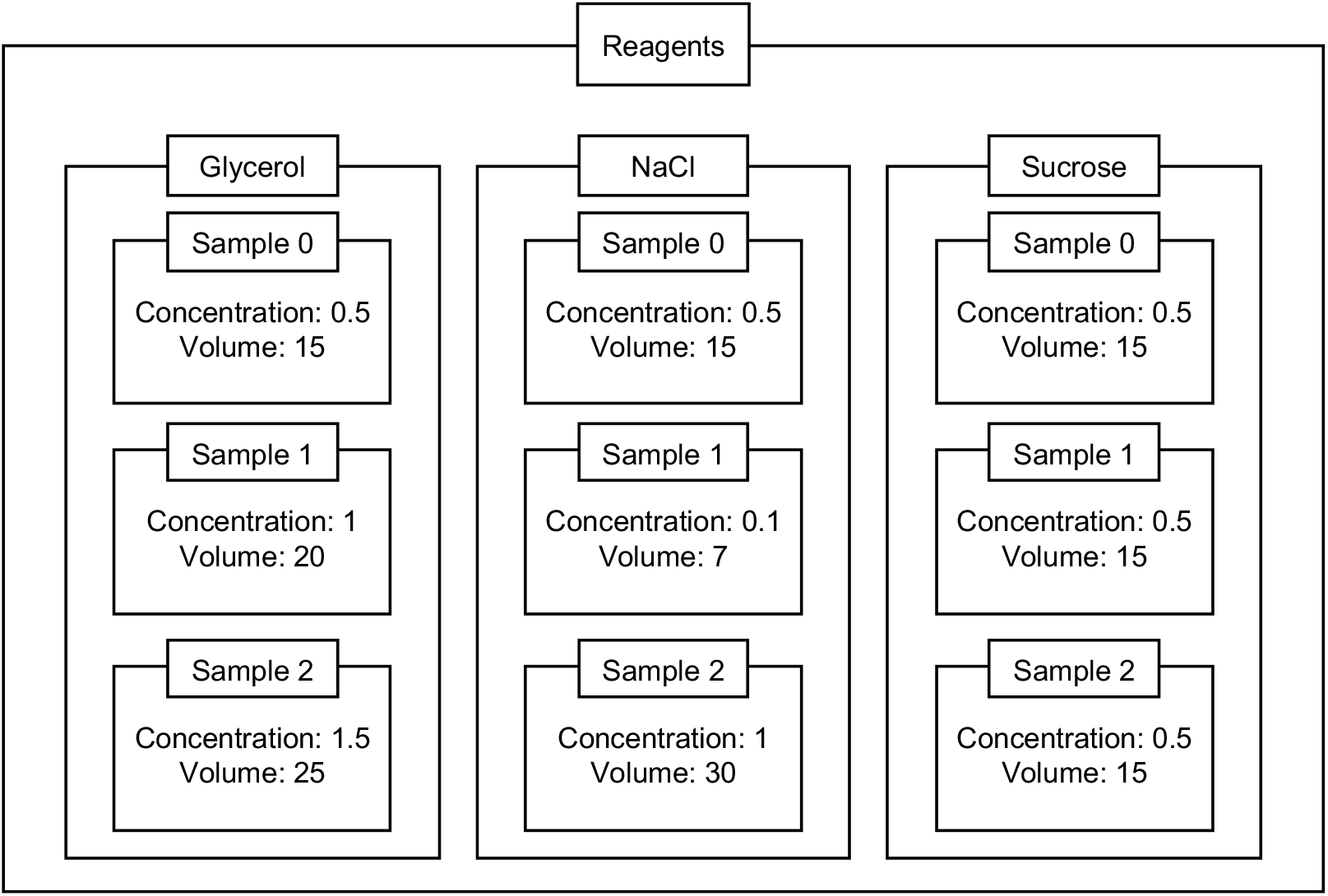
Graphical representation of dictionary data structure within OT-Mation. Values are saved to keys within a dictionary, with values being able to take on most data types available in Python. The Reagents dictionary contains each reagent as a key, the value saved to each reagent key is a nested dictionary where the keys are sample identifiers with values stored as another dictionary. This thrice nested dictionary contains keys matching attributes of interest for the reagent in a given sample such as Concentration as the key and ‘1’ as the value.

After running OT-Mation, a Python output script is generated in the same working directory as the OT-Mation script. Every time the OT-Mation script is run, a new output script will overwrite the existing output script. The output script can be uploaded to the Opentrons companion application, which reads the script and generates instructions for the robot to carry out the experiment. Copies of output script can be created and shared with other researchers allowing users to share and reproduce experiments.

## Conclusion

OT-Mation improves accessibility and efficiency of automated liquid handling for multifactorial experiments using the OT-2 robotic system. By enabling the generation of bespoke Python scripts from user-defined CSV files, OT-Mation bridges the gap between the flexibility of code-based robotics control and the user-friendliness required for broader adoption in laboratory settings. As this tool simplifies the programming process, it also allows researchers with limited coding experience to leverage the full capabilities of the OT-2. As a result, OT-Mation both increases accessibility of advanced liquid handling automation and enhances the reproducibility and accuracy of complex multifactorial experiments.

## METHODS

OT-Mation is implemented in Python 3.9. Testing of the script was performed on an OpenTrons OT-2 Liquid Handling platform.

## ASSOCIATED CONTENT

## Data availability statement

Code, documentation, examples and installation instructions can be found at: https://github.com/tphoward/OT-Mation_Repo.

## AUTHOR INFORMATION

### Author contributions

AL and TPH designed OT-Mation, AL wrote OT-Mation, AL and KC implemented and tested OT-Mation, AL, KC and TPH wrote the manuscript, CH, JT, JS and TPH supervised the research. All authors commented and edited the manuscript.

### Notes

The authors declare no competing financial interest.

## ACKNOWLEDGEMENTS

AL and CH were supported by the Institute for Agri-Food Research Institute (IAFRI). KC was supported by the BBSRC Newcastle-Liverpool-Durham DTP (Doctoral Training Partnership) scheme (reference 2144113). AL and TPH were supported by BB/W01095X/1. For the purpose of open access, the author has applied a Creative Commons Attribution (CC BY) licence to any Author Accepted Manuscript version arising from this submission.

